# High-accuracy mapping of human and viral direct physical protein-protein interactions using the novel computational system AlphaFold-pairs

**DOI:** 10.1101/2023.08.29.555151

**Authors:** Christian Poitras, Felix Lamontagne, Nathalie Grandvaux, Hao Song, Maxime Pinard, Benoit Coulombe

## Abstract

Protein-protein interactions are central, highly flexible components of regulatory mechanisms in all living cells. Over the years, diverse methods have been developed to map protein-protein interactions. These methods have revealed the organization of protein complexes and networks in numerous cells and conditions. However, these methods are also time consuming, costly and sensitive to various experimental artifacts. To avoid these caveats, we have taken advantage of the AlphaFold-Multimer software, which succeeded in predicting the structure of many protein complexes. We designed a relatively simple algorithm based on assessing the physical proximity of a test protein with other AlphaFold structures. Using this method, named AlphaFold-pairs, we have successfully defined the probability of a protein-protein interaction forming. AlphaFold-pairs was validated using well-defined protein-protein interactions found in the literature and specialized databases. All pairwise interactions forming within the 12-subunit transcription machinery RNA Polymerase II, according to available structures, have been identified. Out of 66 possible interactions (excluding homodimers), 19 specific interactions have been found, and an additional previously unknown interaction has been unveiled. The SARS-CoV-2 surface glycoprotein Spike (or S) was confirmed to interact with high preference with the human ACE2 receptor when compared to other human receptors. Notably, two additional receptors, INSR and FLT4, were found to interact with S. For the first time, we have successfully identified protein-protein interactions that are likely to form within the reassortant Eurasian avian-like (EA) H1N1 swine G4 genotype Influenza A virus, which poses a potential zoonotic threat. Testing G4 proteins against human transcription factors and molecular chaperones (a total of 100 proteins) revealed strong specific interactions between the G4 HA and HSP90B1, the G4 NS and the PAQosome subunit RPAP3, as well as the G4 PA and the POLR2A subunit. We predict that AlphaFold-pairs will revolutionize the study of protein-protein interactions in a large number of healthy and diseased systems in the years to come.

## Introduction

Regulation of all cellular processes, from prokaryotes to eukaryotes, require the participation of protein-protein interactions (PPIs). Some of these interactions are relatively stable and give rise to protein complexes, while others are more transient and reversible [1]. PPIs regulate biochemical processes including DNA transcription, replication and repair, RNA translation and maturation, macromolecule turnover, as well as cellular processes such as the immune response, cell cycle and division. PPIs and the assembly of protein complexes are tightly controlled through diverse mechanisms such as posttranslational modifications and the activity of molecular chaperones [2, 3]. For these reasons, understanding the formation of PPIs in detail is a central aspect of cell function to fully comprehend cell growth and differentiation.

Over the years, several methods have been developed to monitor and characterize PPIs. Co-immunoprecipitation, co-localization, two-hybrid assays, phage display and other techniques can be used both at small scale to assess PPIs between two targeted proteins and at a large scale, when combined with techniques like mass spectrometry, to analyze protein networks comprehensively. Protein affinity purification coupled with mass spectrometry (AP-MS) was largely used in the past decade to decipher PPIs in many different cell types and conditions. More recently, proximity dependent biotin identification (BioID and derivatives) was used to map PPIs within their native environment, such as organelles and membranes. Unfortunately, applying these methods is time consuming, expensive and sensitive to experimental artifacts [4]. Moreover, implementing these techniques can pose challenges, particularly when working with samples that possess biohazardous properties. Several groups have sought to use bioinformatic methods to define protein interactomes. However, up to this point, these efforts yielded limited effectiveness [5].

In 2021, the UK company DeepMind, now associated with Alphabet (Google), revealed the development of a neural network termed AlphaFold that performed extremely well in the challenging 14th Critical Assessment of protein Structure Prediction (CASP14), demonstrating accuracy competitive with experimental structures in a majority of cases and greatly outperforming other computational methods [6]. Since publication of the original AlphaFold paper, numerous articles have been published where the AlphaFold network was utilized. As of today, the AlphaFold database contains approximately 200 M structures. Subsequently, a novel iteration of AlphaFold, known as AlphaFold-Multimer, was developed to more effectively determine the structure of protein complexes composed of multiple subunits [7].

In this paper, we describe a new computational method, named AlphaFold-pairs, that takes advantage of the AlphaFold-Multimer software capabilities to identify binary PPIs. Our system uses the physical distance allowed between two proteins in AlphaFold-Multimer. AlphaFold-pairs was validated using two well-defined systems: the 12-subunit enzyme RNA polymerase II (Pol II), and the interaction of the SARS-CoV-2 Spike glycoprotein with human cell surface receptors. Using the AlphaFold-pairs method, we also successfully elucidated the interactions among viral proteins that comprise the reassortant Eurasian avian-like (EA) H1N1 swine G4 genotype Influenza A virus (IAV) variant, a zoonotic respiratory pathogen considered to pose a significant pandemic risk by the CDC and the WHO [8]. In addition, we also identified host interactors that associate with the G4 proteins.

## Experimental section

### Design of AlphaFold-pairs

The usual way to use AlphaFold-pairs is to generate a list of FASTA files from two lists of proteins (baits and targets) by invoking the script “fasta-pairs --baits baits.fasta –targets targets.fasta -u -i --output fasta_pairs”. The script will create a FASTA file for each bait-target combination. Then AlphaFold-multimer is invoked for each FASTA file using the Nextflow pipeline with the command “nextflow run alphafold-pairs.nf --fasta ‘fasta_pairs/*.fasta’”. The number of amino acid (a.a.) pairs that are distanced by 6 Angstroms or less from the “ranked_0.pdb” file created by AlphaFold-multimer can be obtained by using the script “interaction-score -w ranked_0.pdb” or by using the script “multi-interaction-score –w **/ranked_0.pdb” for multiple “ranked_0.pdb” files. This count is normalized by the proteins molecular weight to reduce the false-positive interactions provoked by a greater amount of potential non-specific interacting a.a . The output of “multi-interaction-score” can be converted to a matrix using the script “score-matrix”. The source code is available on GitHub at this URL: https://github.com/benoitcoulombelab/alphafold-pairs.

Protein pairs were analyzed in AlphaFold-multimer and the number of a.a. pairs that are distanced by 6 angstroms and below are charted. This number is normalized according to the size of the proteins. The Z-score of normalized scores (output) was assessed using Perseus (Version 1.6.17.0) when producing heatmaps [9]. The final number of a.a. pairs is used as a score for the interaction. A schematic representation of AlphaFold-pairs is provided in Figure 1.

**Figure 1:**
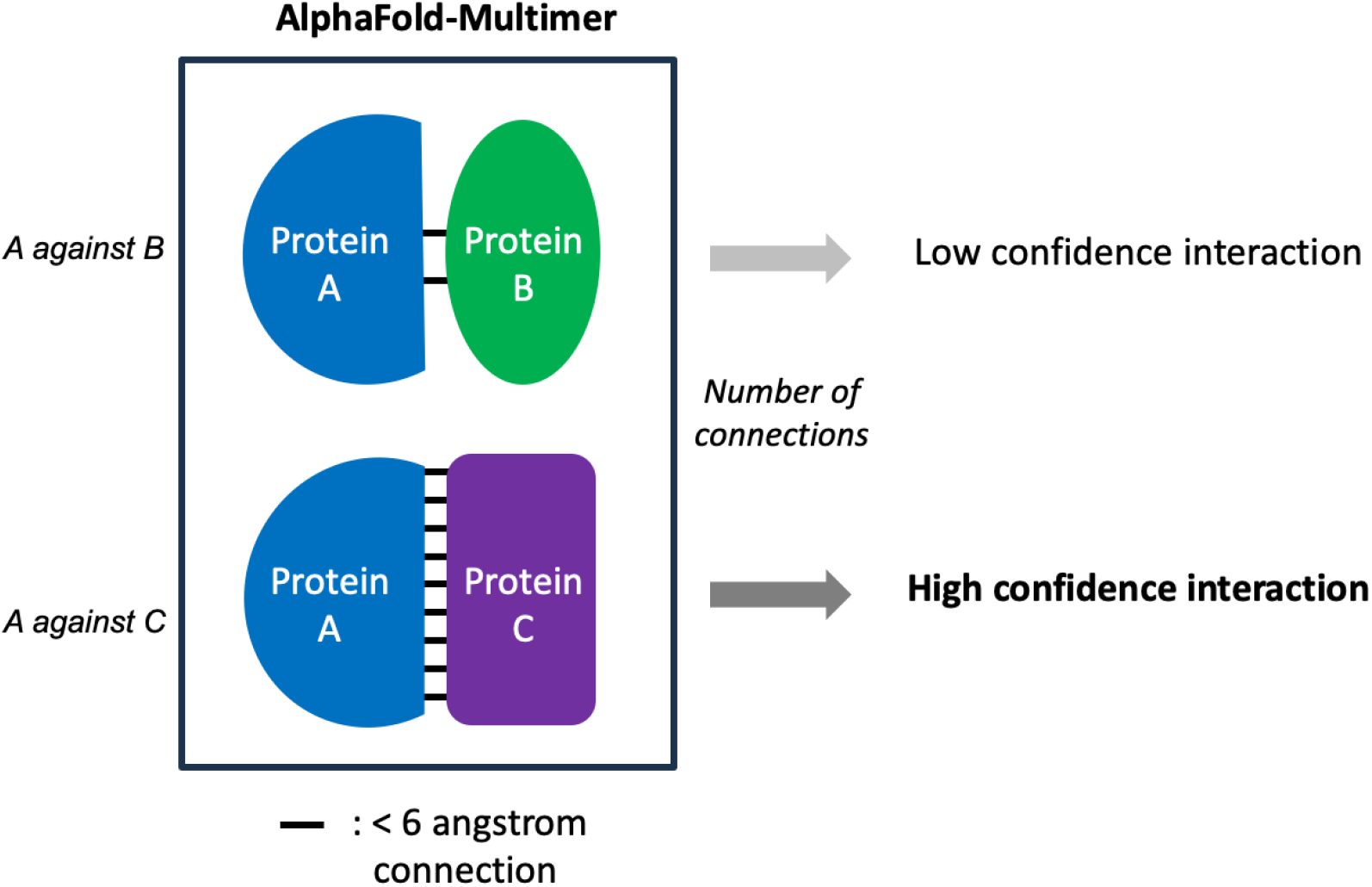
Method overview. In the cartoon, protein A (bait) is compared to the target proteins B and C. In each case, a.a. that are distanced by 6 angstrom or less are monitored and used to define an interaction score. The higher the number of 6 angstrom interactions, the higher the confidence of the interaction.

## Results

### RNA polymerase II subunit pairwise interaction

To assess the capability of AlphaFold-pairs to detect PPIs, we first tested pairwise interactions among subunits of the human Pol II. Pol II is a 12-subunit enzyme that synthesizes mRNA and some other snRNA in all eukaryotic cells. The crystallographic structure of Pol II has been determined at different stages of the transcription reaction using diverse methods [10, 11]. If we disregard homodimeric interactions, a total of 66 interactions are possible and have been tested. In Figure 2 (right), the black ovals are strategically positioned to indicate the subunit contacts observed during structure determination experiment of the enzyme as reviewed by Wild and Cramer in 2012 [12]. AlphaFold-pairs was employed to evaluate every potential pairwise interaction. The thickness of the lines added to Figure 2, corresponds to the confidence level of each interaction. The data indicates that all 19 interactions identified in the structure were detected as high confidence interactions by AlphaFold-pairs. On the other hand, the 47 interactions that were not observed in the structure exhibited low confidence scores, with the exception of the POLR2D-POLR2L interaction, which received a high confidence score. One possibility is that this interaction is formed transiently during the biogenesis or the breakdown of the enzyme. To our knowledge, this interaction has not been reported before. Together, these results validate the effectiveness and specificity of AlphaFold-pairs in accurately identifying binary interactions within a human multi-protein complex.

**Figure 2:**
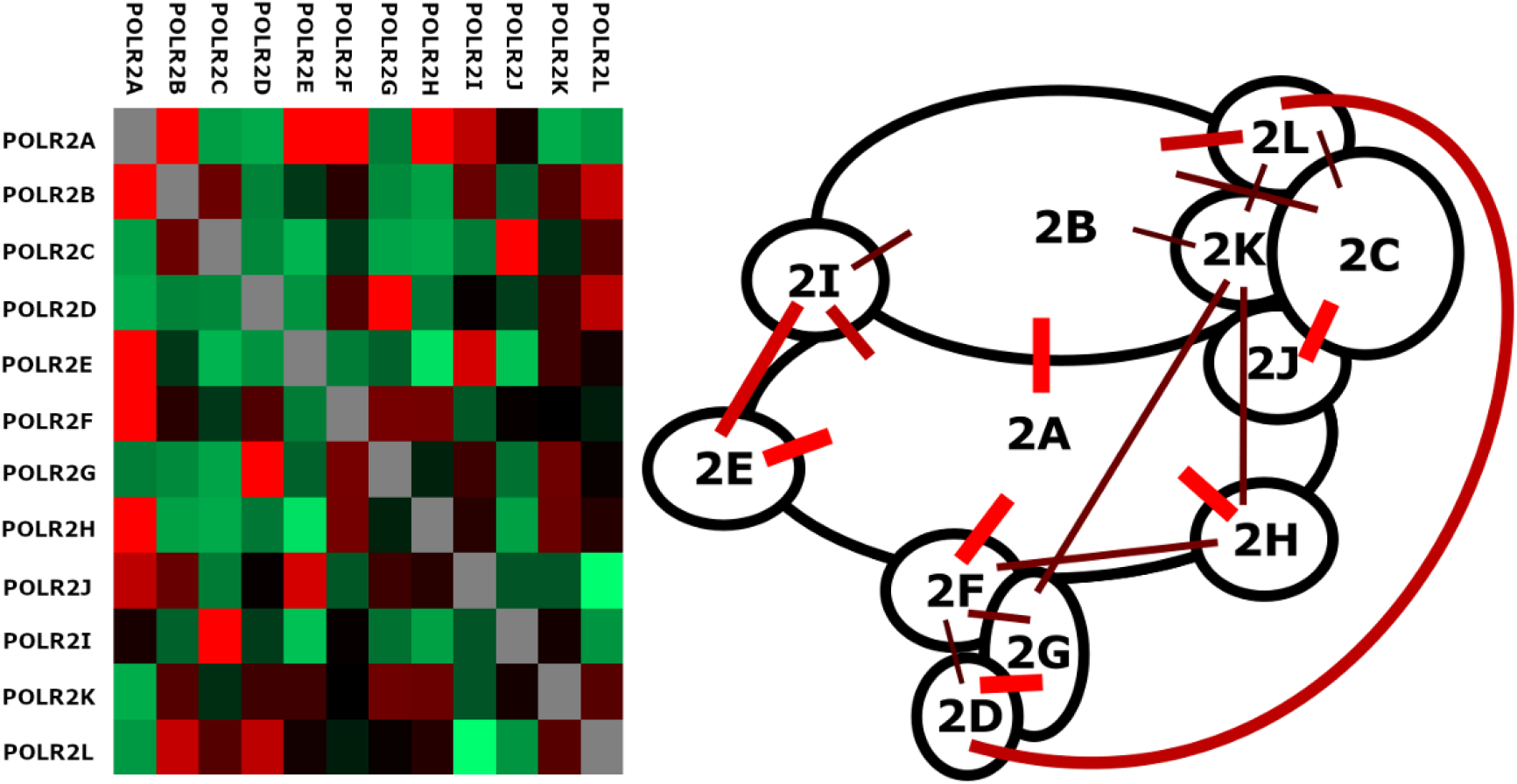
AlphaFold-pairs has been developed to exploit the software AlphaFold-Multimer to systematically identify binary PPIs. The figure shows (left) a heatmap summarizing the interactions between the multiple Pol II subunits, light red being the high confidence interactions. The cartoon (right) represents the structural model of Pol II from the Cramer laboratory with subunits represented using black ovals (12 subunits, POLR2A to POLR2L) [12]. Each possible subunit pair was submitted individually to the AlphaFold-pairs algorithm. Bars represent interaction scores and the color (red to dark red) matches the bar’s thickness. A red bar means a Zscore of 2 or higher while dark read means a Zscore of 0.6. Interactions with scores below 0.5 are not shown. All 19 interactions observed in the structure have high confidence scores.

### SARS-CoV-2 Spike-hACE2 interaction

As part of our validation process, we assessed the pairwise interactions involving the surface glycoprotein Spike of the SARS-CoV-2 virus. Previous studies have demonstrated that Spike specifically binds the human ACE2 receptor (hACE2) [13], facilitating viral recognition and cell entry. In this analysis, we arbitrarily selected 9 receptors of various sizes in addition to hACE2. When compared to the interactions with other cell surface receptors, Spike exhibited a high confidence score for the interaction with hACE2. Again, this experiment demonstrates the efficiency and specificity of AlphaFold-pairs. Unexpectedly, two other receptors, INSR and FLT4, also showed high scores. Recent reports have suggested that individuals who have contracted COVID-19 may have an increased risk of developing diabetes [14]. The INSR-Spike interaction is a possible cause of this condition (see Discussion).

### EA H1N1 swine G4 IAV protein interactions

The reassortant Eurasian avian-like (EA) H1N1 swine IAV G4 is a newly identified reassortant that has emerged in pigs in China and has become predominant since 2016 [15]. The G4 reassortant has HA and NA segments of EA origin, NS gene of triple-reassortant (TR) origin while PB2, PB1, PA, NP and M segments originate from the 2009 pandemic H1N1 virus [15]. In a paper published in 2020 by the Liu laboratory, they reported that the G4 IAV poses a zoonotic threat and has the potential to cause a pandemic [15]. It preferentially binds to human-like sialic acid-alpha-2,6-galactose receptors and is able to infect humans, as evidenced by serological test on swine production workers [15]. The G4 IAV efficiently replicates in human airway epithelial cells and exhibits pathogenicity and transmissibility in ferrets, leading to concerns regarding its potential to cause severe respiratory illnesses in humans [15]. The lack of immunity to this virus among the human population is particularly worrisome and, currently, there is no vaccine available for its prevention or small molecules that can block viral propagation. Using AlphaFold-pairs, we evaluated the intra-virus PPIs within the fourteen ORFs of G4 IAV out of seventeen IAV ORF known [16]. A total of five pairs were identified including the surface glycoproteins HA and NA, the RNA import/export regulation NP and NS, and the three RNA polymerase subunits PA and PB1, PB1 and PB2, PB1 and PB2. It should be noted that interactions obtained with only one member of the pairs are considered hypothetical. Notably, each protein was found to participate in at least one interaction with another subunit, which is expected in the case of tightly interconnected components.

We also tested interactions between G4 proteins and selected host factors. For this experiment, we focused on two groups of human proteins according to GO terms: protein folding chaperone (GO: 0044183) and general transcription factors (GO: 0140223), which include all RNA polymerase subunits. Among these interactions, nine stood out with a score exceeding 14 (see Figure 5). RPAP3 is a component of the HSP90 co-chaperone complex known as the PAQosome [17] while POLR2A is one of the major subunit of the Pol II complex essential for all mRNA transcription. Proteins from other GO terms group like protein transport, hydrolase activity, RNA binding, organelle membrane, immune response, and cell death have been extensively reported.

## Discussion

PPIs play a key role in the regulation of cell processes. Here, we present a computational approach utilizing the AlphaFold-Multimer software to monitor potential PPIs (Figure 1). Through the implementation of the AlphaFold-pairs, we successfully identified interactions in validation experiments, exemplified by the interaction between Pol II subunits (Figure 2) and the binding of the surface glycoprotein Spike from the SARS-CoV-2 virus with hACE2 (Figure 3). This data provides compelling evidence for the efficacy of this method in elucidating PPIs.

**Figure 3:**
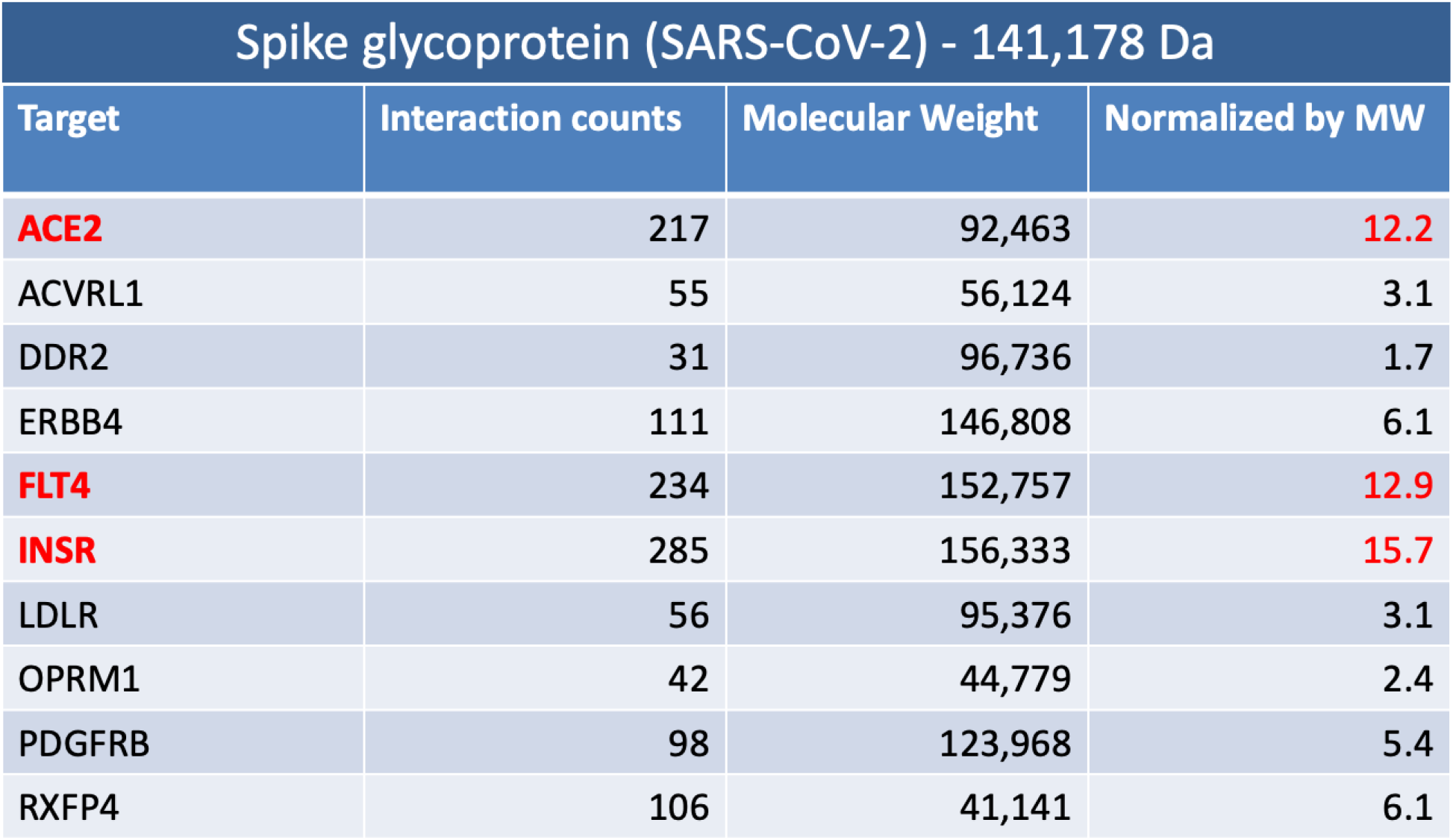
The SARS-CoV-2 S glycopretein was assessed for interaction with a group of human surface receptors of various sizes. The data shows a preference for the ACE2 interaction, as reported previously using other methods. The INSR and FLT4 proteins are also high confidence interactions according to AlphaFold-pairs.

**Figure 4:**
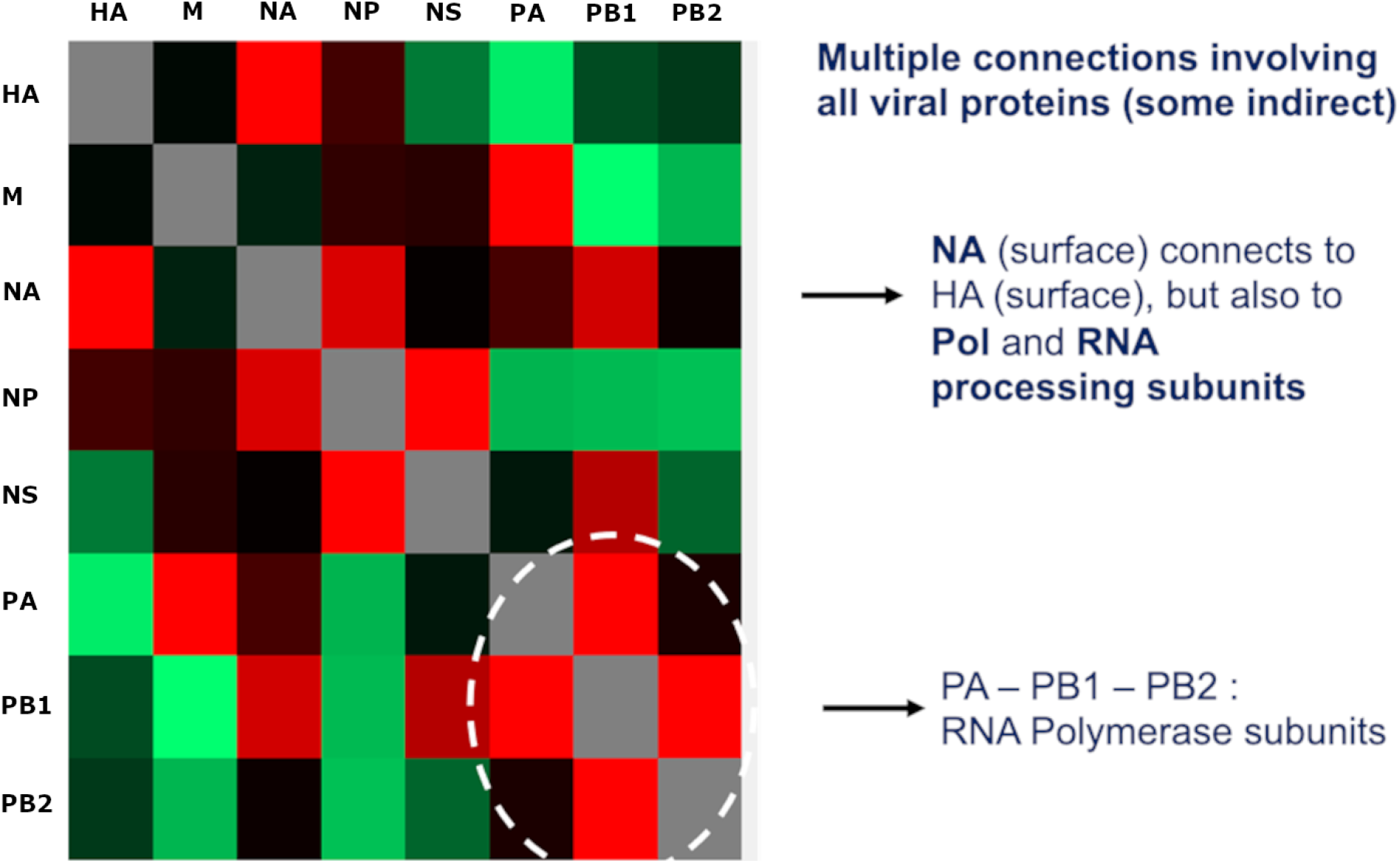
Interactions obtained with the fourteen ORFs of G4 IAV are shown. PB2: Polymerase subunit; mRNA cap recognition ; PB1: Polymerase subunit ; RNA elongation and endonuclease activity ; PA: Polymerase subunit; protease activity ; HA : Surface glycoprotein; major antigen, receptor binding and fusion activities ; NP: RNA binding protein; nuclear import regulation ; NA : Surface glycoprotein; sialidase activity and virus release ; M1: Matrix protein; vRNP interaction, RNA nuclear export regulation and viral budding ; M2 : Ion channel; virus uncoating and assembly ; NS1: Non-structural; Interferon antagonist protein and regulation of host gene expression ; NEP/NS2: Non-structural; nuclear export of RNA.

**Figure 5.**
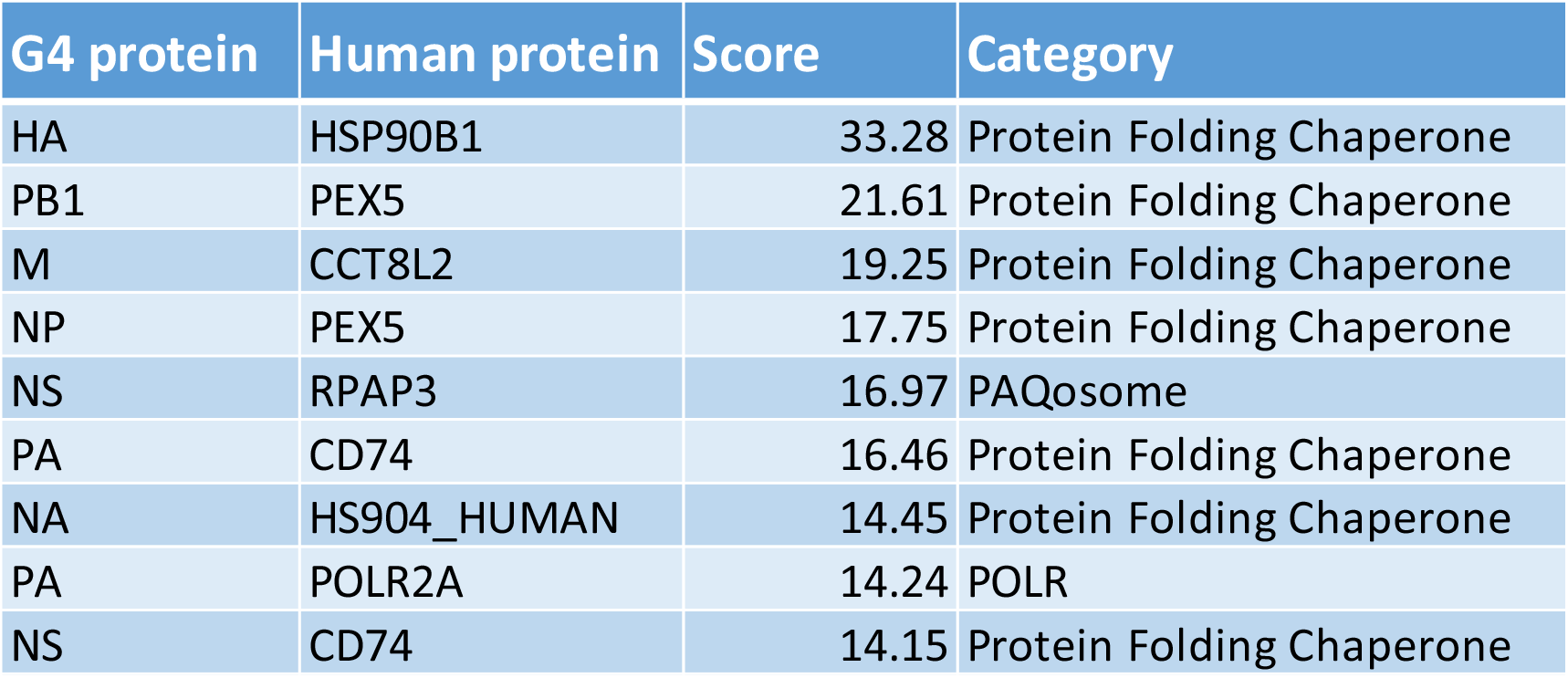
List of the protein-protein interactions between G4 IAV proteins and human transcription factors and molecular chaperones. The highest score is obtained between HA and HSP90B1. Interestingly, a strong interaction is also obtained between NS and the HSP90 co-chaperone RPAP3, a core subunit of the PAQosome.

In addition, by utilizing AlphaFold-pairs, we not only confirmed previously known interactions, but also uncovered novel interactions within both systems. Notably, the identification of the INSR-Spike interaction presents a compelling mechanism through which the SARS-CoV-2 virus may potentially induce diabetes in infected patients [14], as it may interfere with the regular binding of insulin to its receptor. However, it is important to note that experimental validation is necessary to confirm this discovery.

We also exploited the G4 IAV virus as a means to investigate viral PPIs, without the need for virus manipulation. Considering the risk and constrains associated with handling highly pathogenic viruses, AlphaFold-pairs offers a safe approach to unveil viral and viral-host PPIs sincethese organisms require a host to replicate and continue their propagation. Interacting with cell machinery is required to transcribe the viral genome into proteins [18] and the produced proteins can blocked normal cell functions [19]. In this paper, we report on the PPIs that establish connections between viral proteins, as well as interactions with host factors. While the exact function of the interaction between viral proteins with host factors remains unknown, they are potential candidates for antiviral research [20]. These findings are likely to significantly advance our understanding of virus replication and infection processes. Our results so far indicate that AlphaFold-pairs represents a novel and highly valuable tool for studying PPIs.

## Acknowledgements

We thank members of our laboratories for helpful discussions. BC is funded by the Bell-Bombardier Chair of Excellence at the IRCM and the Ministry of Economy and Innovation, Government of Québec. NG received funds from the Fondation du CHUM. FL is recipient of a studentship from the Réseau en Santé Respiratoire du Québec. NG and FL are members of the Réseau en Santé Respiratoire du Québec and Réseau Québécois COVID – Pandémie. NG is a member of the CoVarr-Net network. This research was possible with the support provided by Calcul Quebec (www.calculquebec.ca/) and Compute Canada (www.computecanada.ca).

